# Neural language processing across time, space, frequency and age: MEG-MVPA classification of intertrial phase coherence

**DOI:** 10.1101/2021.10.02.462796

**Authors:** Mads Jensen, Rasha Hyder, Britta U. Westner, Andreas Højlund, Yury Shtyrov

**Affiliations:** Research Unit for Robophilosophy and Integrative Social Robotics, Institut for Kultur og Samfund, Aarhus University, Aarhus, Denmark; Heinrich-Heine University Düsseldorf; Radboud University, Donders Institute for Brain, Cognition and Behaviour, Nijmegen, The Netherlands; Department of Linguistics, Cognitive Science and Semiotics Aarhus University; Aarhus University

## Abstract

Language is a key part of human cognition. Whereas many neurocognitive abilities decline with age, for language the picture is much less clear and how exactly language processing changes with aging is still unknown. To investigate this, we employed magnetoencephalography (MEG) and recorded neuromagnetic brain responses to auditory linguistic stimuli in healthy participants of younger and older age using a passive task-free paradigm and a range of different linguistic stimulus contrasts, which enabled us to assess neural language processes at multiple levels (lexical, semantic, morphosyntactic). By using machine learning-based classification algorithms to scrutinise intertrial phase coherence of MEG responses in source space, we found significant differences between younger and older participants across several frequency bands and for all tested processing types, which shows multiple changes in the brain’s neurolinguistic circuits which may be due to both healthy aging in general and compensatory processes in particular.

## Introduction

Language is a key part of the human cognitive inventory, acquired early in life and used until death. As we age, some cognitive abilities decline, which goes in parallel with several physical changes in the brain (Raz et al., 2005). However, people can produce and understand language until death, and aging comes with both negative (such as difficulties in name retrieval) and positive (increased vocabulary) alterations in language functions (Abrams & Farrell, 2011). But exactly how the language function changes with age and what mechanisms ensure its resilience is still poorly understood. Another reason why age-related dynamics in language functions are important is that several neurodegenerative diseases typically linked with older age (such as Alzheimer’s or Parkinson’s disease) lead to a cognitive decline in patients and the cognitive decline is often correlated with age; such pathological changes and healthy ageing effects may be difficult to disentangle. To understand the impact that a neurodegenerative disease has on the cognitive aspects of language processing, we first need to understand the normal aging-related changes. In this research report, we address this question by investigating the impact of age on different types of linguistic information processing in the brain, namely the lexical (word storage), semantic (meaning), and syntactic (linguistic structure parsing) levels of comprehension.

Structurally, age-related brain changes mostly manifest as shrinkage across most of the tissues, including neocortical grey matter, hippocampus, and white matter tracts, all of which include parts of the left-lateralised language network (Davis et al., 2009; Pfefferbaum et al., 2000; Raz et al., 2005). However, these structural changes as such do not lead to a measurable loss in language comprehension (Agarwal et al., 2016), which raises the question of how the language function can functionally resist obvious changes in its neuronal underpinnings. While it is fairly uncontroversial to suggest that some form of compensation must take place, it is important to specify what mechanisms underly such compensatory changes that can keep the functional output the same in spite of neuroanatomical degradation.

Another situation where compensatory processes are known to take place is during successful rehabilitation after brain injuries where core areas of the left-hemispheric language network are damaged. Several lesion studies have reported an increase in the right hemisphere’s activity after damage to the left-hemispheric language network (Blasi et al., 2002; Kielar et al., 2016; Thiel et al., 2006; Tyler, Wright, et al., 2010). This suggests that the right hemisphere can “step in” to help perform language processing if needed (although lesion studies per se do not tell us about what happens in a healthy brain). Similar to these findings, it has also been reported that there is an increase in right-hemispheric activity with normal ageing. For instance, Tyler et al. (2010) argued that, with age, syntactic processing switches from a primarily left-hemispheric frontotemporal system to a bilateral functional language network. Agarwal et al. (2016) reported that the language areas develop more bihemispheric functional connectivity with age, despite neuroanatomical losses. In another recent study, Gertel et al. (2020) have reported a bilateral activation of frontal areas and precuneus in an older group compared to left-lateralised activity in younger individuals in a lexical task. Interestingly, this relative increase in right-hemispheric involvement does not seem to be at the expense of the left hemisphere, but rather reflects a more distributed response. Still, the nature of this “redistribution” of functional activity remains obscure.

Language comprehension is a highly dynamic process unfolding on a millisecond scale, which is best addressed using time-resolved neuroimaging methods, such as electro- or magnetoencephalography (EEG, MEG). Such EEG/MEG studies have most often focussed on event-related potentials/fields (ERPs/ERFs; for a review see e.g. Friederici, 2002). In recent years, however, more studies have examined the neural oscillatory dynamics underpinning normal language processing. For instance, θ-oscillations (∼4-6 Hz) have been shown to play a role in both acoustic processing and sentence parsing (see e.g. Bastiaansen & Hagoort, 2006; Kösem & van Wassenhove, 2017; Luo & Poeppel, 2007). More importantly in the context of the present report, higher-frequency oscillations have been related to different neurolinguistic processes. Bastiaansen and Hagoort (2006) suggested that both β (∼15-25 Hz) and *γ* (>30 Hz) bands are related to the unification of semantic and syntactic information. Towle and colleagues (2008) have, in turn, shown that power in the high *γ* band (70-100 Hz) is elevated when hearing words compared to hearing tones, linking it to lexical processing, while θ and *γ* band activity is involved in processing acoustic dynamics or speech signal (Teng et al., 2017). These and many other studies have led to a more general suggestion that, while low frequencies reflect the analysis of the acoustic features, high-frequency activity in the *γ* range is related to perceived linguistic representations, and the entirety of speech processing is underpinned by the interplay between low- and high-frequency neural oscillations (Kösem and van Wassenhove, (2017). In sum, there is substantial evidence that the brain handles different linguistic information by engaging oscillatory activity at different frequencies, and multiple neurophysiological frequency bands are involved in fully functional language processing networks in an interactive manner.

Generally, patterns of neural oscillations (also those not related to linguistic processing) have been shown to change with age. For instance, using simple visual stimuli, Gaetz and colleagues (2012) showed that the peak frequency for gamma activity decreases with age. Furthermore, Ziegler and colleagues (2010) reported an increase in β-band activity with age using both real data and simulations in the primary somatosensory cortex with tactile stimuli. Another study (Schafer et al., 2014) used spectral analysis of neural oscillations in resting state data to show that there is an increase in inter-regional amplitude correlations with age, largest in α and β frequency bands. Jointly, these and other results (obtained with non-linguistic paradigms, e.g. Gaetz et al., 2010, 2012; Herrmann et al., 2011) show an influence of age on oscillatory brain activity, even though the direction of those changes (i.e., increase or decrease) vary across individual frequency bands, while the mechanisms and the functional role of these changes still remain to be specified.

Given the link between oscillations and language processing on the one hand, and a change in oscillatory brain dynamics with age on the other, one way to investigate the changes in language processing occurring with age could be to address language-related neural oscillations in different age groups. This was the objective of the present study, where we used MEG to record automatic brain responses to different linguistic contrasts and subsequently applied source modelling and machine learning techniques to disentangle oscillatory signatures of different linguistic processing levels in younger and older participants.

One common issue when addressing neural processes in individuals with different cognitive or neurological status is the risk of potential confounds related to differences between tasks or groups in attentional levels, cognitive resources available for the task or motor vigilance for providing responses (Gansonre et al., 2018; Hyder et al., 2020). Thus, in order to investigate differences in language processing between younger and older persons in the absence of task/attention demands and related confounds, we devised a passive task that does not require any stimulus-related behavioural responses from the participants and that could be conducted without relying on any attentional resources. Passive tasks have been used successfully in many studies to show language processing without attention (e.g. Hyder et al., 2020, 2021; Naatanen et al., 2007; Pulvermüller et al., 2006). In our task, the participants were exposed to spoken words played over headphones while watching a film without sound in the MEG scanner, similar to previous studies on automatic language comprehension processes that do not require attention on the auditory input or any responses at all (Naatanen et al., 2007; Pulvermuller & Shtyrov, 2006; Shtyrov, 2010). This allowed us to investigate age-specific differences based on automatic responses to the same stimuli without attentional or motor requirements confounding the results. To assess different linguistic processing, we used spoken stimuli that diverged lexically (words/pseudo-words), semantically (action-related/abstract), or morphosyntactically (grammatically correct/ungrammatical).

We have previously shown that oscillations linked to automatic processing of spoken words can be used to classify different types of linguistic information in the input (M. Jensen et al., 2019), where especially the phase of the oscillatory activity in several frequency bands allowed for successful decoding of language features. We found that this classification was most successful for lexical processing across several distinct *γ* sub-bands, for semantic processing – in the α and β bands, and for syntactic processing – in the low *γ* band. Thus, to investigate age-related processing differences, we focused on analysing intertrial phase coherence (ITPC) in five canonical frequency bands (α, β, and low, medium, and high γ), with two different age groups consisting of healthy younger and older participants. Further, for neuroanatomical localisation of the cortical oscillatory dynamics, we opted to compute the ITPC in source space using a cortically constrained linearly-constrained minimum variance (LCMV) beamformer (Van Veen et al., 1997), calculated using individual brain-based boundary element models. We chose multivariate pattern analysis (MVPA) as our statistical approach since it can handle large amounts of data compared to traditional frequentist statistics and allows for unbiased data driven analysis without having to a priori specify either locations, time points, or frequency bands of interest. Combined, this approach allowed us to decode different language properties over time in the individual frequency bands, and to compare activity across the two groups of younger and older participants.

## Methods

### Participants

MEG data were acquired for two groups of participants, all right-handed (assessed according to Oldfield, 1971) healthy native Danish speakers with normal hearing and no record of neurological impairments: seventeen healthy young participants (age range 18–27 years, mean age 22.95 years, 12 females) and sixteen older participants (age range 51-75 years, mean age 64 years, 11 females). All participants gave written consent and received remuneration for their participation. The experiment was approved by the Central Denmark Region Committees on Health Research Ethics and was conducted according to the principles of the Helsinki Declaration.

### Stimuli

To address a range of different neurolinguistic processes at the lexical, semantic, and syntactic levels, we chose stimulus items which could enable us to contrast a combination of different linguistic phenomena while controlling for acoustic features (see Fig. 1 for examples of the stimuli used). To this end, we followed a previously suggested strategy (Gansonre et al., 2018) and selected a set of spoken Danish-language stimuli which (i) belonged to different lexical and semantic categories (action-related verb, abstract verb, object-related noun and meaningless pseudo-word), (ii) were close in terms of phonology so we could compare them directly with minimal acoustic/phonetic confounds, and (iii) could be modified morpho-syntactically in the exact same way and nonetheless exhibit different syntactic properties (i.e., grammatically correct vs. incorrect) such that we could test the very same contrasts in different linguistic contexts in a counterbalanced fashion.

**Figure 1:**
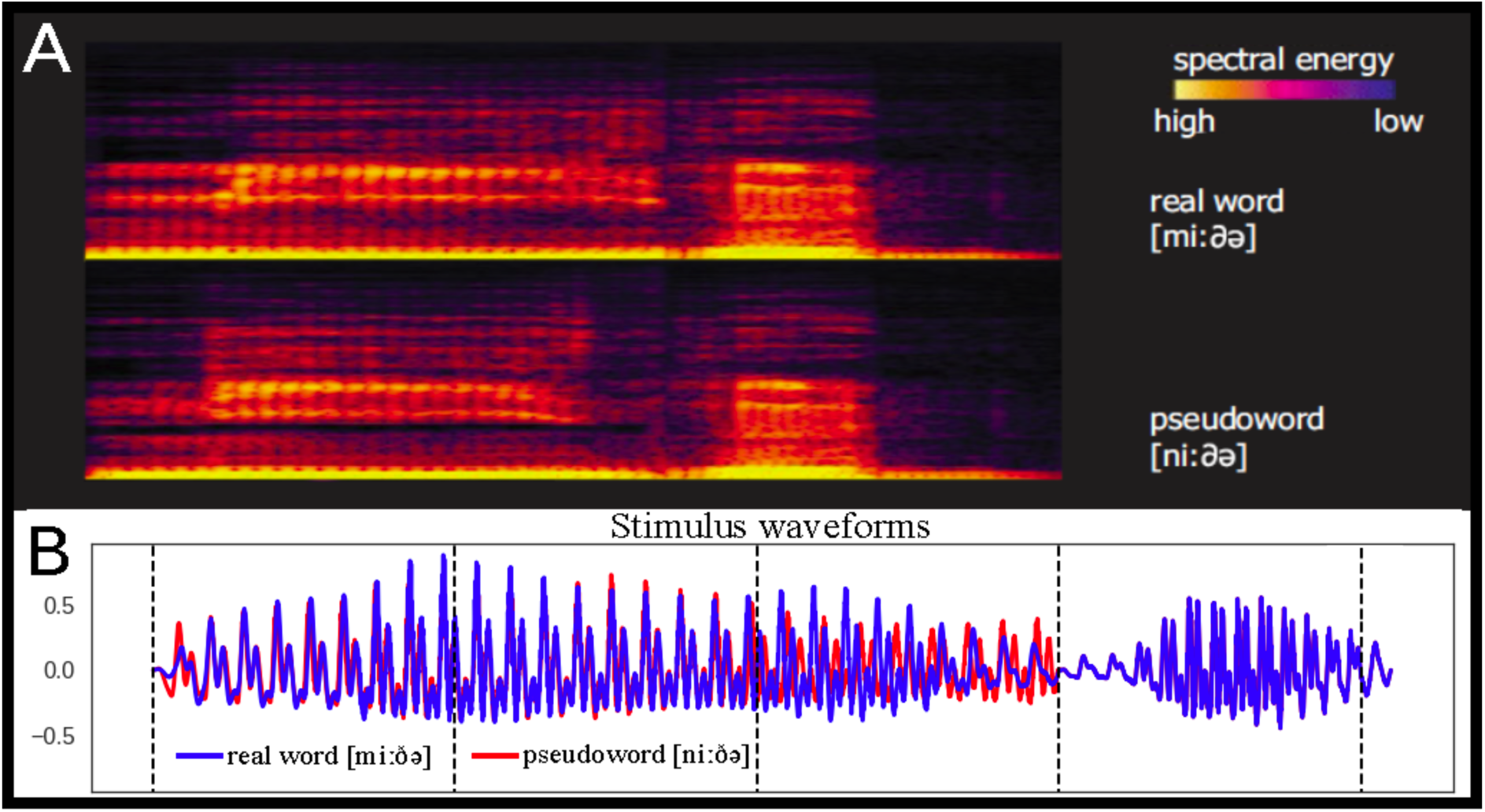
Example stimulus. Adapted from Jensen et al. (2019). ***A,*** Examples of spectrograms of spoken stimuli used in the experiment. ***B***, Examples of waveforms plotted on top of each other.

This led to a choice of four main base stimuli: bide ([biːðə], *to bite*), gide ([giːðə], *to bother*), mide ([miːðə], *a mite*), *nide ([niːðə], \**pseudo-word*). These were presented as such. Note that these words have identical CVCV phonological structure and only differ in the first consonant. The second syllable [ðə], which allows recognition of the lexical items, was the same across all items. To ensure that the full recognition of each word form in the restricted experimental context is only possible at the second syllable, we also included, in a 1:1 ratio with all other stimuli, all four first syllables in isolation: [biː], [giː], [miː] and [niː]. These served as fillers to ensure identical acoustic divergence points across the four types of stimuli, to which we time-locking the brain activity, and were thus not analysed as such.

The above stimulus quadruplet provided us with a way to address both lexical and semantic contrasts. By estimating the brain activity elicited by the same word-final syllable [ðə], we could compare, on the one hand, word vs. meaningless pseudo-word activation, putatively indicating lexical access, which we expected to be reflected in an automatic activation of the core left temporo-frontal language system (Tyler & Marslen-Wilson, 2008). On the other hand, by comparing action vs. non-action items, we could address semantically-specific aspects of these activations. Previous EEG, MEG and fMRI research has indicated automatic involvement of the brain’s motor system in the comprehension of action-related verbs (Pulvermüller, 2005; Pulvermüller & Fadiga, 2010); we therefore expected more pronounced centro-frontal activity for the action verb *bide*, but not for the concrete noun *mide*.

We produced, based on the forms above, further stimuli that included a balanced morphosyntactic contrast. We took advantage of Danish morphology and the fact that the morphemes *-(e)t* and *-(e)n* can be used to express the past participle of verbs and definiteness on common nouns. Hence, we compared the inflected items based on their syntactic congruence or incongruence, e.g. *-n* in mide*n* vs. *gide*n*, and *-t* in gide*t* vs. *mide*t* (where * indicates a violation of the stem/affix syntactic agreement). Note that all of these pairs have identical codas (*t/n*) that lead to grammatical/morphosyntactic violation in a counterbalanced fashion: each of them is correct in combination with one but not the other stem. These were presented, in equal proportion along with the other stimuli above, to make sure syntactic properties were only recognised at the very last consonant. To balance for these acoustic modifications, we also included similar items based on other forms (*bide[n/t] and *nide[n/t], all meaningless), which were used to make a balanced design, but not analysed as such.

The stimuli were made based on a digital recording of a male native speaker of Danish in an anechoic chamber (recording bandwidth: 44k Hz, 16bit, stereo). The first and second syllables of four CVCV stimuli were recorded independently, in order to avoid possible coarticulation effects, and cross-spliced together, such that the second syllables were physically identical across all items. The second syllable commenced at 300 ms after the onset of the first one, and this was the earliest time (the so-called disambiguation point) when any lexical or semantic effects could be expected in the MEG data.

To produce the morphosyntactic items, ending in [t] or [n], we cross-spliced recordings of these two morphemes onto the four main stems in order to obtain words either violating or respecting rules of Danish morphology such that the exact same phonemes completed syntactic or asyntactic forms in a counterbalanced fashion. These morphemes became distinct at 408 ms after the word onset, and this was therefore the earliest time any morphosyntactic contrasts could affect the brain responses.

The sounds were matched for loudness, with a 1.93 dB drop between the first and the second syllables so that our stimuli sounded as natural as possible and they were normalised to have identical power (measured as root-mean-square, RMS). All sound editing was done using Adobe Audition CS6 software (Adobe Inc., San Jose, CA).

To summarise, the stimulus set included four CV syllables, four CVCV stems, four CVCV+[t] and four CVCV+[n] forms, all strictly controlled for phonological and acoustic properties. These were combined in a pseudorandom fashion in a single auditory sequence ensuring that the stimuli’s lexical, semantic and syntactic properties were available at stringently defined times.

### Procedure

The MEG recording was conducted in an electromagnetically shielded and acoustically attenuated room (Vacuumschmelze Gmbh, Hanau, Germany). During the recordings, participants were placed in supine position and instructed to focus on watching a silent film (displayed using a bespoke mirror system) and to pay no attention to the sounds. The auditory stimuli were controlled using Neurobehavioral Systems Presentation v16 (www.neurobs.com) and presented through in-ear-tubes (Etymotic ER-30) binaurally at 50 dB above individual auditory threshold.

All sixteen stimuli were presented equiprobably in a single data acquisition session inter- mixed continuously with 100 pseudo-random repetitions of each stimulus resulting in 1600 epochs in total. The total recording time was 28 minutes.

MEG data were acquired with an Elekta Neuromag Triux MEG (MEGIN Oy, Helsinki, Finland) with 102 magnetometers and 204 planar gradiometers. For eye movement and heartbeat artefact detection, 2 bipolar electrooculogram (EOG) and 1 bipolar electrocardiogram (ECG) recordings were taken. Cardinal landmarks and additional head points were digitised using a Polhemus FASTRAK setup (Polhemus, Vermont, USA). Data were recorded at 1000 Hz, a high-pass filter of 0.1 Hz and low pass of 330 Hz were applied online. Head position and head movements were continuously tracked using four Head Position Indicator (HPI) coils. The participants were lying still on a non-magnetic patient bed, with their head as close to the top of the helmet as possible, the MEG dewar being in supine position, and the display mirror fixed above the head.

### MEG data preprocessing

All data were preprocessed using the MNE-Python open source software package, v. 0.19 (Gramfort et al., 2013). First, the raw continuous data were bandpass filtered from 1 to 95 Hz and downsampled to 500 Hz and then epoched into single-trial epochs of 1000-ms duration, starting 100 ms before and ending 900 ms after stimulus onset. Bad channels were detected automatically and interpolated using an automatic approach (Jas et al., 2017). Epochs with excessive bad channels were discarded and outlier trials removed. For the young group there was on average 23.24 bad channels (median: 32, SD: 10.93) and 12.9 (SD: 18.66) bad epochs. For the older group there was on average 2.62 bad channels (median: 2, SD: 1.59) and 8.83 (SD: 78.69) bad epochs. Finally, cleaned epoched data were bandpass-filtered into five frequency bands (Dalal et al., 2011): α (8-12 Hz), β (13-30 Hz), low-γ (30-45 Hz), medium-γ (55-70 Hz) and high-γ (70-90 Hz).

### Source reconstruction

For each participant, a T1 structural magnetic-resonance image (MRI) was obtained using a Siemens Prisma 3T MRI scanner (Siemens Healthcare Gmbh, Germany). The images were segmented in order to create surfaces for the inner skull using SimNIBS software (Thielscher et al., 2015). For each subject, an individual 1-layer boundary element model (BEM) and individual forward model were calculated. A common template grey matter surface was created by averaging all the study participants using the Freesurfer software (Dale et al., 1999).

Source reconstruction was carried out based on planar gradiometer data using a unit-noise-gain scalar LCMV beamformer (Van Veen et al., 1997) and a Hilbert transformation-based beamforming technique (Westner, 2017; Westner & Dalal, 2017). For the adaptive filter, source orientation was optimised by using the orientation of maximum signal power; the output value selected was the neural activity index (NAI, Sekihara & Nagarajan, 2008). The strategy of using only gradiometer data was chosen as planar gradiometers are less sensitive to external magnetic interference and have a better signal-to-noise ratio compared to magnetometers; furthermore, combining channel types is not trivial due to magnetometers and gradiometers producing values of different scales and units (Hari et al., 1988). First, for each frequency band of interest (α, β, low-γ, medium-γ, and high-γ), the epochs were bandpass-filtered for each subject *without* subtracting the evoked signal from the single trials as we were interested in investigating the complete information in the responses time-locked to the auditory stimuli. Second, an adaptive filter was created by combining responses to all the stimuli in the paradigm using a 3-layer BEM and a cortically constrained source space. The adaptive filter was computed using a data covariance matrix based on all time points of the bandpass-filtered epochs; the covariance matrix was not regularised prior to inversion. Third, after the adaptive filters were created, the single-trial epoched data were Hilbert-transformed and the adaptive filter was applied to the complex Hilbert-transformed data providing a source reconstruction for each single trial. Last, we calculated the inter-trial phase coherence (ITPC) of the obtained single-trial source space data. This was done for each time point and in each source space location independently using the equation below:

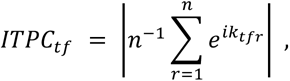

where *n* is the number of trials, *e^ik^* provides a complex polar representation of the phase angle *k* on trial *r* for the time-frequency point *tf* where frequency is the frequency band (for a review Cohen, 2014 see Chapter 19, esp. pp. 244-245). This resulted in a single ITPC time course for each frequency band and for source point for each subject.

After the source space ITPC data were calculated for each subject, the individual data were morphed onto the common template surface (5124 vertices). Finally, the data were smoothed with a 10-ms rolling windowed mean in the temporal dimension for each source independently.

### Multivariate pattern analysis

For each participant, the morphed ITPC time series per source point was extracted based on the contrast in question. Common to all the multivariate pattern analyses (MVPA) was the strategy of classification over time, i.e., independently for each time point a classifier pipeline was applied providing a classification score for this time point, which then, taken together, created a classification score over time. We used the entire cortical source space as input for the classification at each time point, providing 5124 features per time point.

The classifier pipeline was constructed in MNE-Python using scikit-learn (Pedregosa et al., 2011) and composed of three steps: First, the features were standardised (z-scored), which was done across all vertices in the source space at each time point independently using the training set and then applied to the test set. Second, the best 1024 features were selected using F-values (implemented in scikit-learn’s SelectKBest algorithm)^1^. Last, a logistic regression (C=1, L-BFGS solver, number of max iterations = 1000) was used to classify the two contrasts, and the receiver operating characteristic area under the curve (ROC-AUC) was used as the classification score. This pipeline was applied across subjects, meaning that we obtained one classification score over time for all participants and only looked for effects that could hold across the subjects in the tested group.

In order to avoid overfitting of the MVPA models, we used cross-validation where we created a training set of all the subjects except one and then used the left-out subject for testing the model (called leave-one-subject-out; see Varoquaux et al., 2017 for details and strategies for cross-validating brain imaging data). This was done iteratively across subjects, until all subjects had been used for as a testing set once. In the end, we obtained one ROC-AUC score for each participant.

This pipeline was then repeated for both groups and each frequency band within each linguistic process independently resulting in an average ROC-AUC score for each band and contrast for each group. To assess the statistical significance of the classification, we opted for a two-step approach. In the first step, we calculated the threshold for a significant binomial test based on the number of participants and stimuli. In the second step, we corrected for multiple comparisons using a cluster-based approach; based on previous literature (Edmonds & Krumbholz, 2014; Hagoort & Brown, 2000; Wang, Jensen, et al., 2012), we deployed a threshold of 10 ms such that only clusters lasting 10 ms or longer were considered of interest.

## Results

### Lexical contrast

In the lexical contrast, we found that we could successfully classify real words vs. pseudo- words for both the younger and the older groups. The times and frequency bands of successful classification, however, varied for the two groups. For the young group we obtained significant classification in the α, medium-ɣ, and high-ɣ bands in seven different time-clusters (see Table 1); in the older group, we obtained significant classification in the β and medium-ɣ bands with two different time-clusters. The earliest time point of decoding was much earlier for the young group in the high-ɣ already at 48 ms after the divergence point, compared to 266 ms after the divergence point for the older group in the β band (see Table 1 and Figs. 2 & 3 for an overview). The highest ROC-AUC score or the young group was 81.58% (SD: 5.00) in the medium-ɣ band in a time-cluster from 74-100 ms after the divergence point. For the older group, it was 78.12% (SD: 3.44) in the medium-ɣ band in a time-cluster from 276-294 ms after the divergence point.

**Figure 2:**
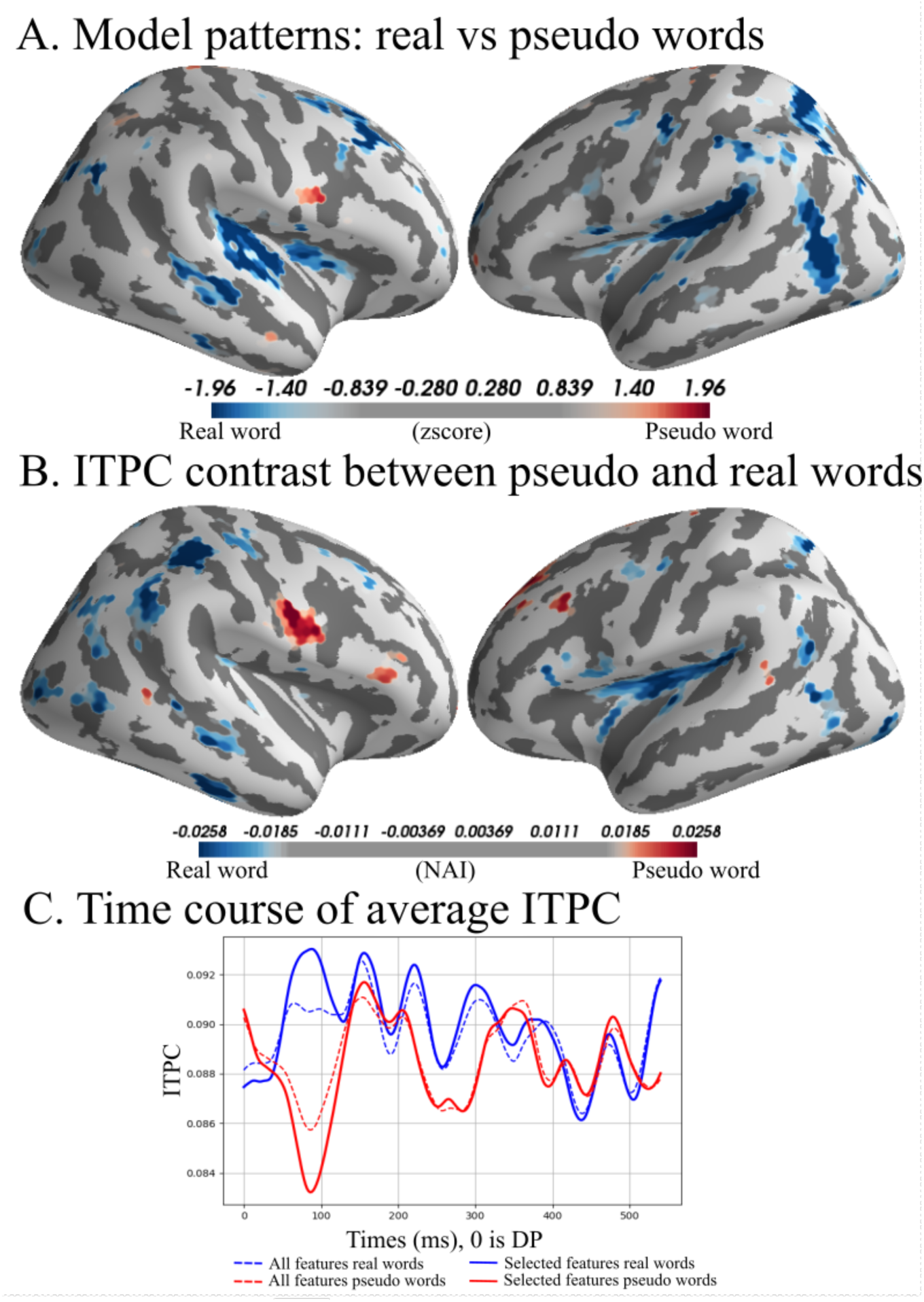
Lexical condition, young group, medium-ɣ, 74 - 100 ms after divergence point. ***A***, Model patterns: in order to interpret the coefficients of the machine learning model we use model patterns (for details, see Haufe et al., 2014). We show the top and bottom 5% of the patterns in the medium-ɣ band, from 74 to 100 ms for the young group. Blue colours are areas of activation that predict real words and red are areas used to predict pseudo words. ***B***, Average top and bottom 5% of ITPC difference; blue colours indicate higher ITPC for real words and yellow/red colours indicate higher ITPC for pseudo word for the medium-γ band from 74 to 100 ms. ***C***, Average ITPC over time; solid lines are the average of the selected features, dashed lines are the average of all vertices in the source space. Time 0 is the divergence point, when stimuli could be distinguished from the available acoustic information.

**Figure 3:**
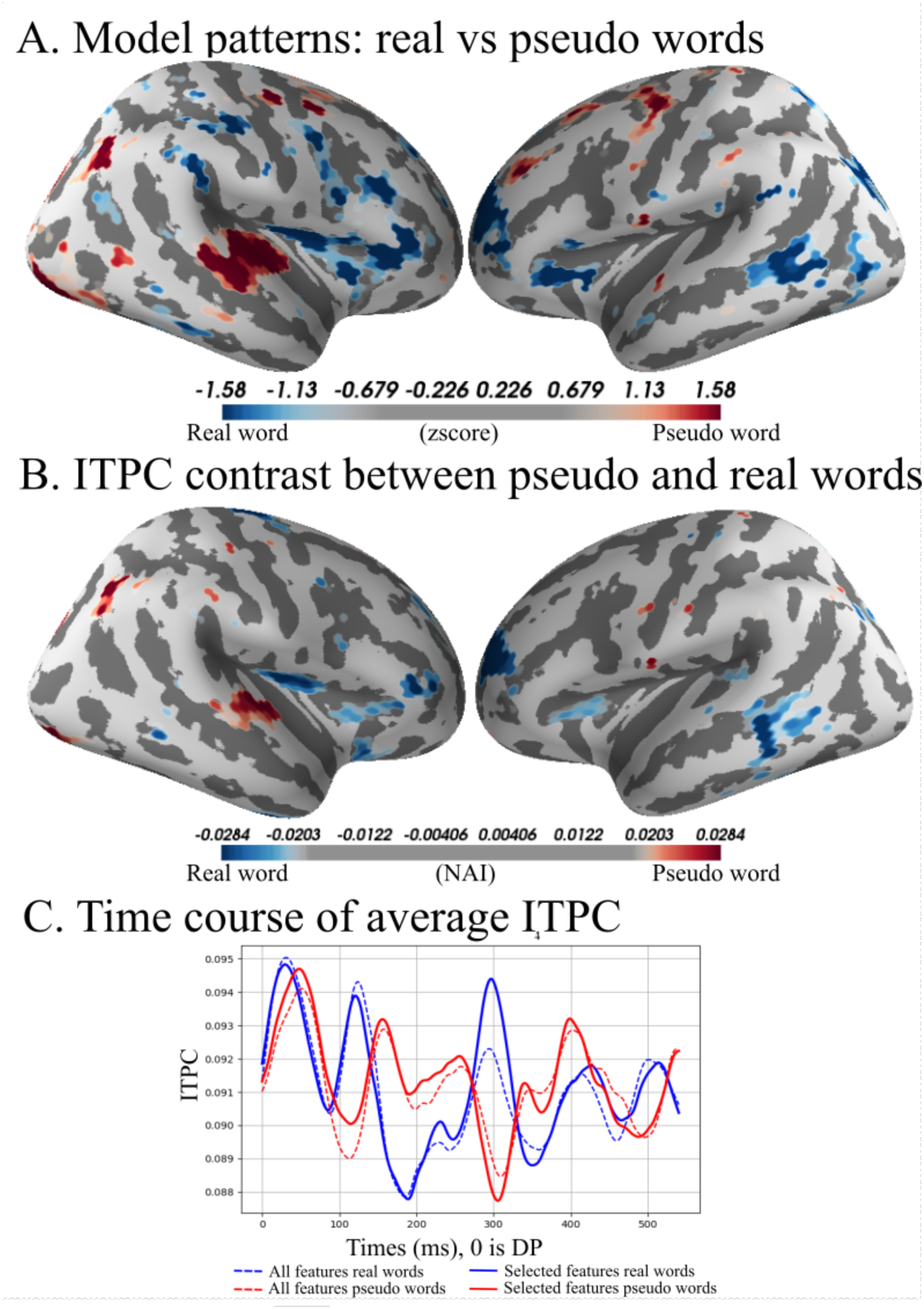
Lexical condition, older group, medium-ɣ, 276 - 294 ms after divergence point. ***A***, Model patterns (see also Fig. 2 legend): top and bottom 5% of the patterns in the medium- ɣ band, from 276 to 294 ms in the older group. Blue colours are areas that predict real words and yellow/red are areas that predict pseudo words. ***B***, Average top and bottom 5% of ITPC difference, blue colours indicating higher ITPC for action verb and yellow/red indicating higher ITPC for object noun from 276 to 294 ms in the older group. ***C***, Average ITPC over time, solid lines are the average of the selected features, dashed lines are the average of all vertices in the source space. Time 0 is the divergence point, when stimuli could be recognized from the available acoustic information.

**Table 1:**
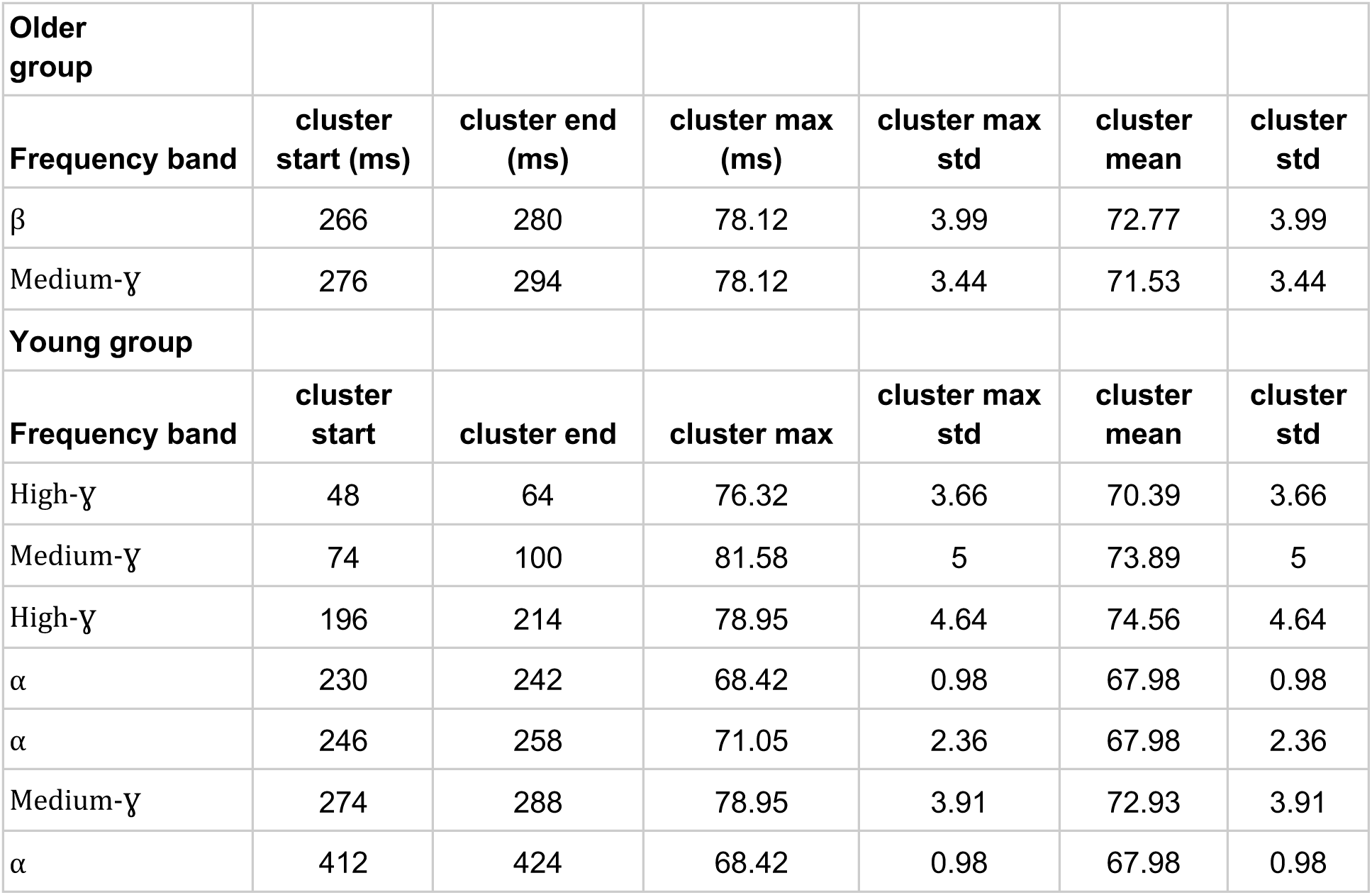
Lexical condition. Cluster start is the start time of the cluster from DP and *cluster end* is the end time of the cluster from DP. C*luster max* is the highest ROC-AUC score within the cluster. The *cluster max std* is the standard deviation of cross-validation folds for the max ROC-AUC score. *Cluster mean* is the mean ROC-AUC score of the cluster. The *cluster std* is the standard deviation of the cluster mean across cross-validation folds.

Inspection of the left-hemispheric patterns from the time-cluster with the peak ROC-AUC score for the young group showed a pattern of activity predicting the real word in Brodmann areas (BA) 39, 40, 41, 44 as well as BA 1, 4, 6 and 7. In the right hemisphere we found patterns predicting the real word in BAs 21-22, 39 as well as 6 and 8, with a small cluster in BA 6 predicting the pseudo-word. For the other, less prominent time-clusters, we found mostly bilateral patterns that predicted both the real word and the pseudo-word.

For the older group the patterns from the cluster with the peak ROC-AUC score in the left hemisphere showed smaller patterns of activity, which could predict the real word, in BA 21- 22, as well as 3 and 6. There were also smaller patterns predicting the pseudo-word in BA 38, 39, 46 as well as 2, 3, and 11. In the right hemisphere, we found patterns predicting the real word in BA 6, 8, 19 and BA 43. Predicting the pseudo-word were patterns in BA 22, 39, 40, 46 as well as 2,1,3, and 11.

### Semantic contrast

For the semantic contrast, we could successfully classify action verb vs. object noun for both the younger and the older groups. Again, the times and frequency bands of successful classification varied for the two groups. For the younger group, we found significant classification in the β and medium-ɣ bands in two different time-clusters, while in the older group five different time-clusters were found in the α and medium-ɣ bands. The first time point of decoding for the young group was in the β band at 142 ms after the divergence point, compared to 154 ms after divergence point for the older group in the α band (see Table 2 and Figs. 4 & 5 for an overview). The highest ROC-AUC score for the young group was 78.95% (SD: 4.73) in the β band in the time-cluster from 142-158 ms after the divergence point. For the older group, it was 79.41% (SD: 1.96) in the α band in the 192-226 ms cluster.

**Figure 4:**
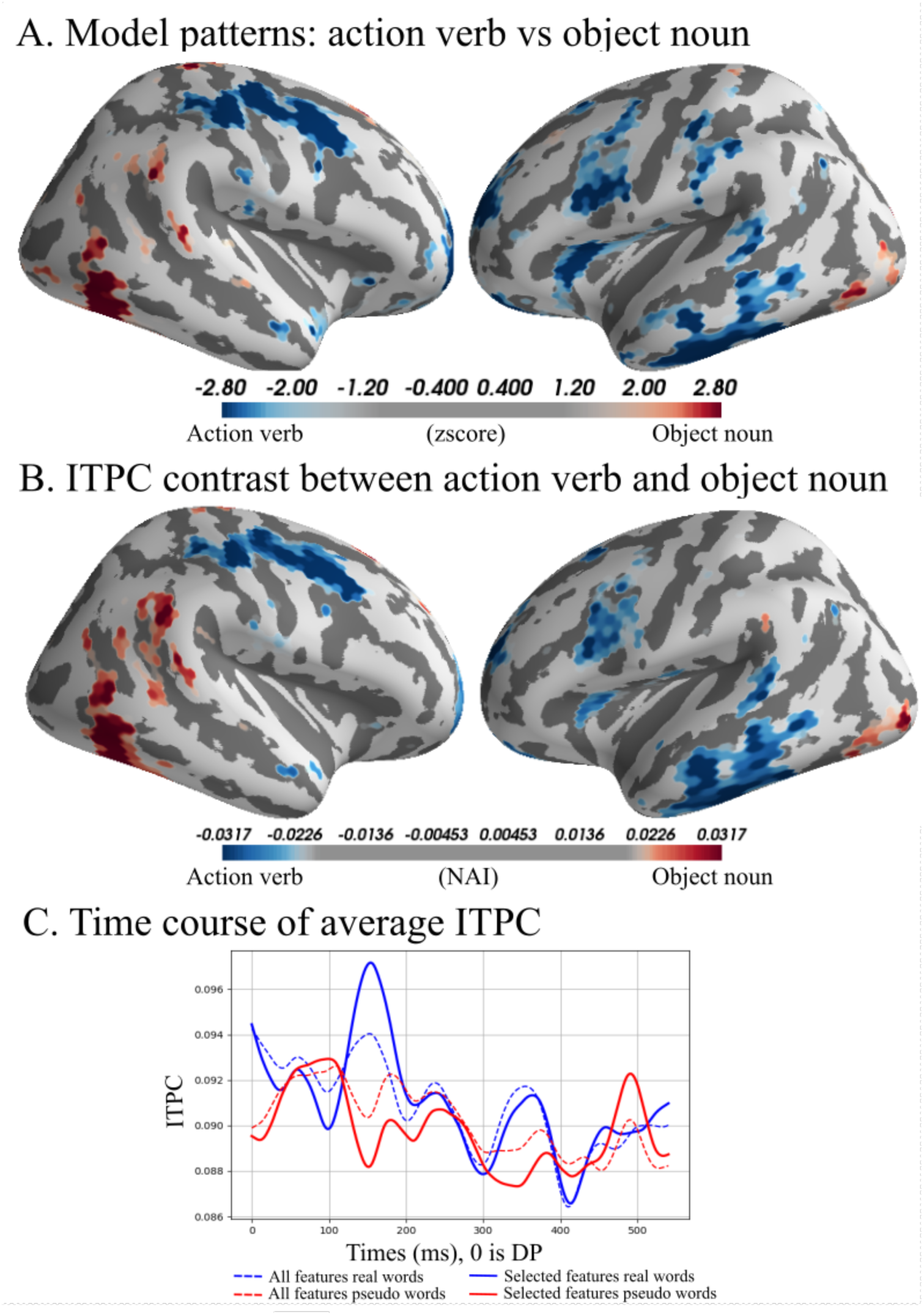
Semantic condition, young group, β, 142 - 158 ms after divergence point. ***A***, Model patterns (see also Fig. 2 legend): top and bottom 5% of the patterns in the β band from 142 to 158 ms in the young group. Blue colours are areas used to predict action verb and yellow/red are areas used to predict object noun. ***B***, Average top and bottom 5% of ITPC difference, blue colours showing higher ITPC for action verb and yellow/red indicating higher ITPC for object noun from 142 - 158 ms in the young group. ***C***, Average ITPC over time, solid lines are the average of the selected features, dashed lines are the average of all vertices in the source space. Time 0 is the divergence point, when stimuli could be recognised from the available acoustic information.

**Figure 5:**
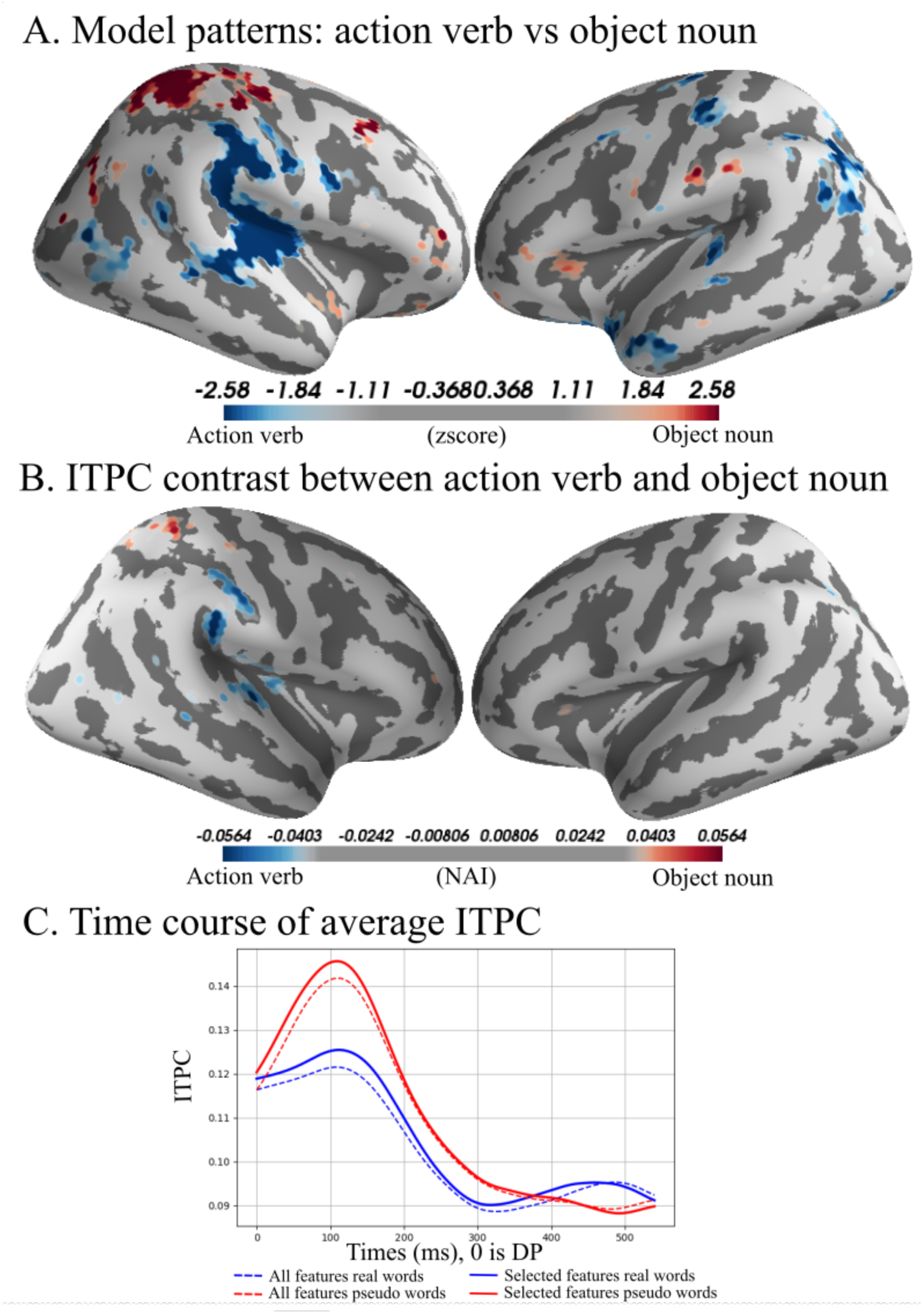
Semantic condition, older group, α, 382 - 414 ms after divergence point. ***A***, Model patterns (see also Fig. 2 legend): top and bottom 5% of the patterns in the α band, from 382 to 414 ms in the older group. Blue colours depict areas used to predict action verb and yellow/red are areas used to predict object noun. ***B***, Average top and bottom 5% of ITPC difference, blue colours showing higher ITPC for action verb and yellow/red indicating higher ITPC for object from 382 to 414 ms in the older group. ***C***, Average ITPC over time, solid lines are the average of the selected features, dashed lines are the average of all vertices in the source space. Time 0 is the divergence point, when stimuli could be recognised from the available acoustic information.

**Table 2:** Semantic condition. Cluster start is the start time of the cluster from DP and *cluster end* is the end time of the cluster from DP. C*luster max* is the highest ROC-AUC score within the cluster. The *cluster max std* is the standard deviation of cross-validation folds for the max ROC-AUC score. *Cluster mean* is the mean ROC-AUC score of the cluster. The *cluster std* is the standard deviation of the cluster mean across cross-validation folds.

|  |  |  |  |  |  |  |
| --- | --- | --- | --- | --- | --- | --- |
| **Elderly group** |  |  |  |  |  |  |
| **band** | **cluster start (ms)** | **cluster end (ms)** | **cluster max (ms)** | **cluster max std** | **cluster mean** | **cluster std** |
| $\alpha$ | 154 | 170 | 73.53 | 1.95 | 71.32 | 1.95 |
| $\alpha$ | 192 | 226 | 76.47 | 2.45 | 73.18 | 2.45 |
| Medium-γ | 256 | 276 | 79.41 | 3.73 | 72.65 | 3.73 |
| $\alpha$ | 382 | 414 | 79.41 | 3.1 | 75.37 | 3.1 |
| Medium-γ | 464 | 476 | 76.47 | 2.77 | 72.55 | 2.77 |
| **Young group** |  |  |  |  |  |  |
| **Band** | **cluster start (ms)** | **cluster end (ms)** | **cluster max (ms)** | **cluster max std** | **cluster mean** | **cluster std** |
| $\beta$ | 142 | 158 | 78.95 | 3.89 | 74.34 | 3.89 |
| Medium-γ | 486 | 498 | 71.05 | 1.96 | 70.18 | 1.96 |

Inspecting the patterns from the time-cluster with the peak ROC-AUC score for the young group of the left hemisphere showed a pattern with activity predicting the action verb in a large set of areas: BA 6, 44, 20, BA 22 and 40, and BA 37. In the right hemisphere, we also found patterns predicting the action verb in sensorimotor areas: BA 2, 4 and 6; the patterns predicting the object noun was, in turn, found in inferior-temporal cortex (fusiform gyrus, BA 37). For the medium-ɣ band in the left hemisphere, we only found a small cluster in BA 22 predicting action verb, and another one in BA 10 predicting object noun. In the right hemisphere, object noun-predicting clusters were located in BA 44, 4, 6.

For the older group, the patterns from the time-cluster with the peak ROC-AUC score in the left hemisphere showed activity predicting the object noun in temporal areas BA 22, 37 and 38. In the right hemisphere, a cluster predicting object noun was in the temporoparietal junction (TPJ; BA 39).

### Morphosyntactic contrast

We could successfully classify correct vs. incorrect morphosyntax for both groups. As with the lexical and semantic contrasts, the times and frequency bands of successful classification varied between the two groups. For the young group, significant classification was achieved in the α and high-ɣ bands in two different time-clusters; in the older group, it was achieved in the β and high-ɣ bands in four different time-clusters. The earliest time point of successful decoding was for the young group in the β band at 346 ms after the divergence point, as opposed to 104 ms in the α band in the older group (see Table 3 and Figs. 6-7). The highest ROC-AUC score for the young group was 65.79% (SD: 2.63) in the α band in a time-cluster from 412-424 ms after the divergence point. For the older group, it was 72.06% (SD: 1.71) in the β band in a time-cluster from 148-158 ms after the divergence point.

**Figure 6:**
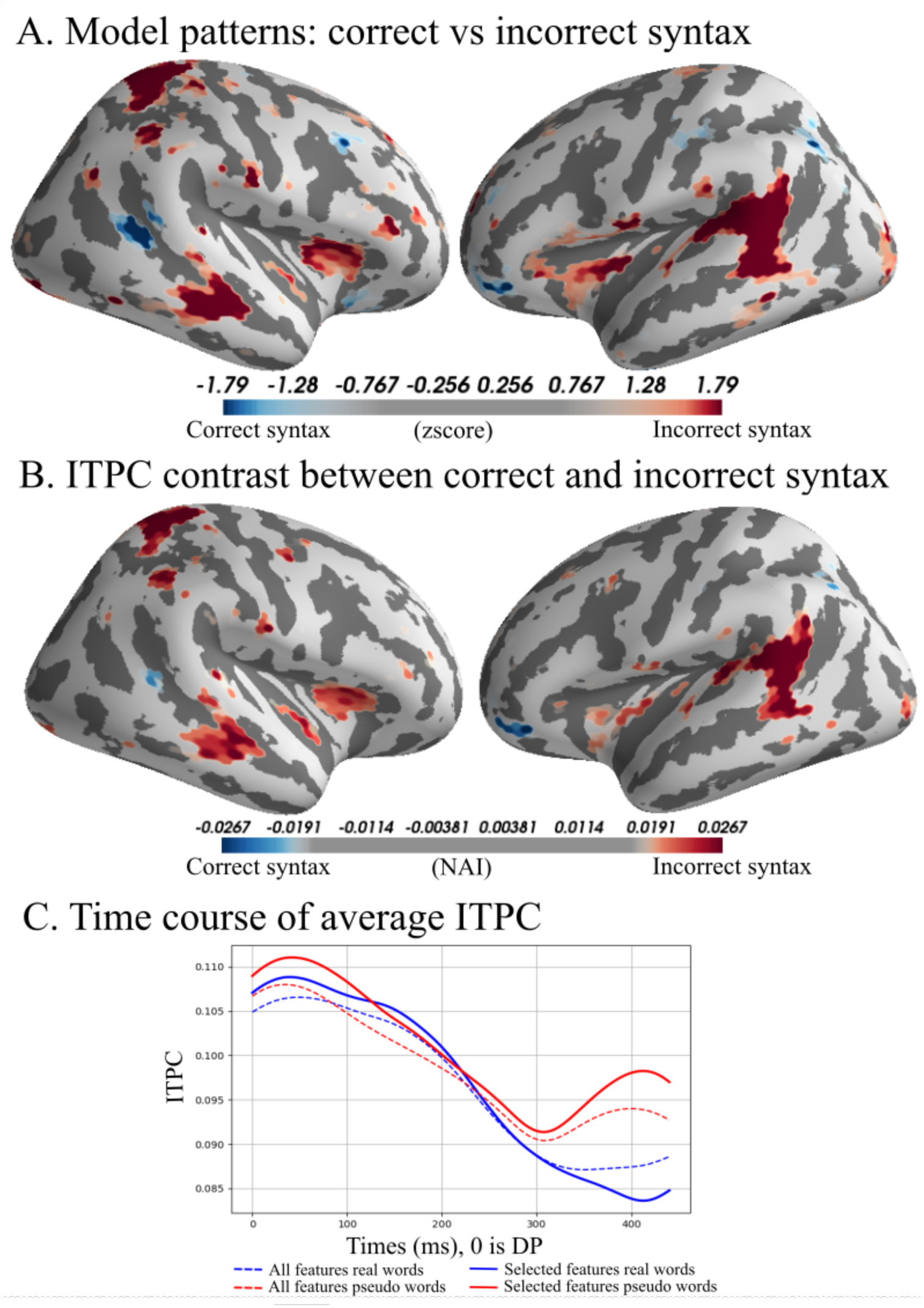
Morphosyntactic condition, young group, α, 412 - 424 ms after divergence point. ***A***, Model patterns (see also Fig. 2 legend): top and bottom 5% of the patterns in the α band, from 412 to 424 ms in the young group. Blue colours are areas used to predict correct syntax and yellow/red are areas used to predict incorrect syntax. ***B***, Average top and bottom 5% of ITPC difference, blue colours showing higher ITPC for correct syntax and yellow/red showing higher ITPC for incorrect syntax from 412 to 424 ms in the older group. ***C***, Average ITPC over time, solid lines are the average of the selected features, dashed lines are the average of all vertices in the source space. Time 0 is the divergence point, when stimuli could be recognised from the available acoustic information.

**Figure 7:**
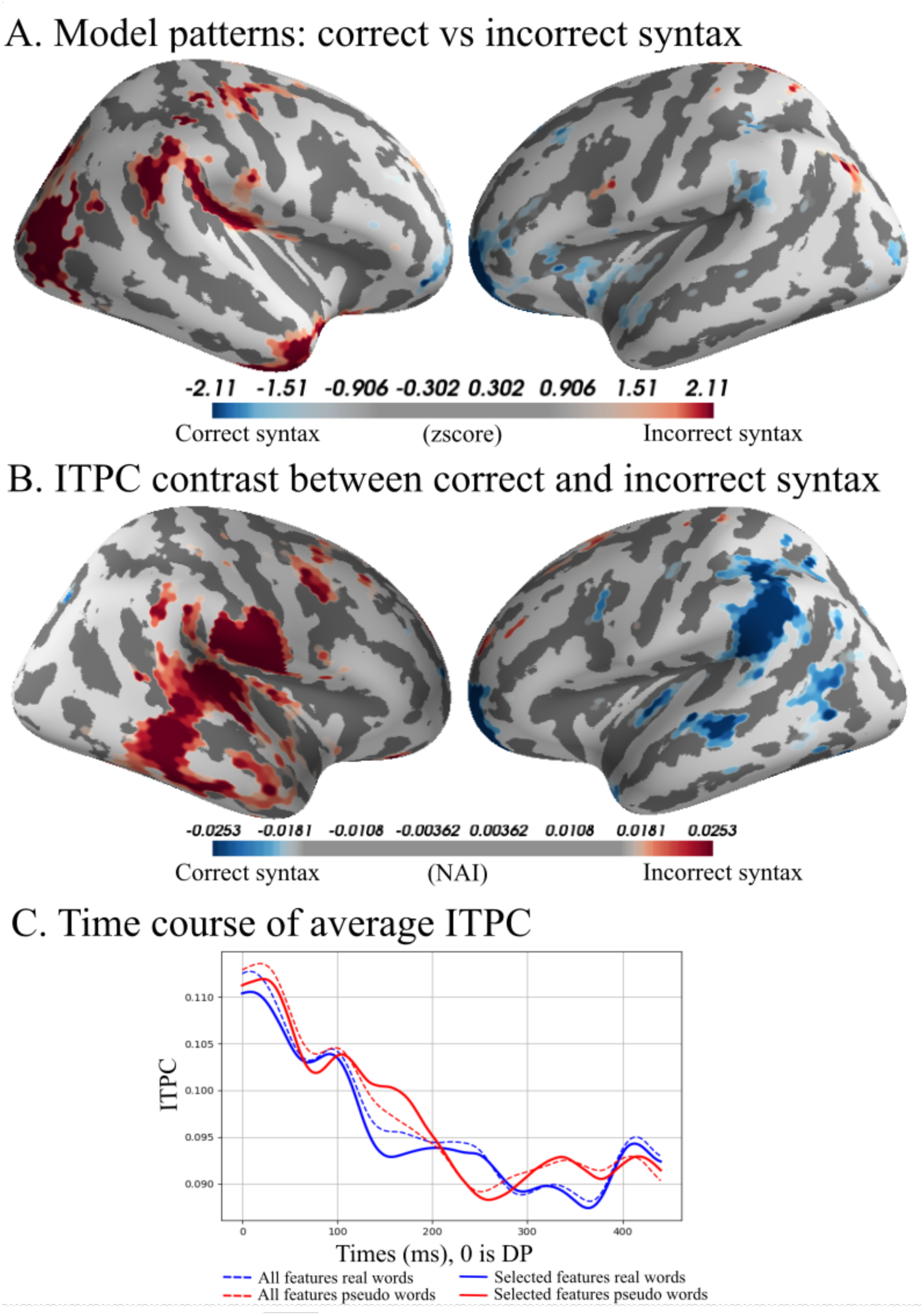
Morphosyntactic condition, older group, β, 148 - 158 ms after divergence point. ***A***, Model patterns (see also Fig. 2 legend): top and bottom 5% of the patterns in the β band, from 148 to 158 ms in the older group. Blue colours are areas used to predict correct syntax and yellow/red are areas used to predict incorrect syntax. ***B***, Average top and bottom 5% of ITPC difference blue colours showing higher ITPC for correct syntax and yellow/red showing higher ITPC for incorrect syntax from 148 to 158 ms in the older group in the older group. ***C***, Average ITPC over time, solid lines are the average of the selected features, dashed lines are the average of all vertices in the source space. Time 0 is the divergence point, when stimuli could be recognised from the available acoustic information.

**Table 3:** Morphosyntactic condition. *Cluster start* is the start time of the cluster from DP and *cluster end* is the end time of the cluster from DP. C*luster max* is the highest ROC-AUC score within the cluster. The *cluster max std* is the standard deviation of cross-validation folds for the max ROC-AUC score. *Cluster mean* is the mean ROC-AUC score of the cluster. The *cluster std* is the standard deviation of the cluster mean across cross-validation folds.

|  |  |  |  |  |  |  |
| --- | --- | --- | --- | --- | --- | --- |
| **Elderly group** |  |  |  |  |  |  |
| **band** | **cluster start (ms)** | **cluster end (ms)** | **cluster max (ms)** | **cluster max std** | **cluster mean** | **cluster std** |
| $\beta$ | 104 | 124 | 72.06 | 4.45 | 67.94 | 4.45 |
| $\beta$ | 148 | 158 | 72.06 | 1.71 | 68.82 | 1.71 |
| High-γ | 256 | 270 | 70.59 | 4.37 | 62.61 | 4.37 |
| $\beta$ | 430 | 440 | 67.65 | 3.14 | 65 | 3.14 |
| **Young group** |  |  |  |  |  |  |
| **band** | **cluster start (ms)** | **cluster end (ms)** | **cluster max (ms)** | **cluster max std** | **cluster mean** | **cluster std** |
| High-γ | 346 | 356 | 64.47 | 10.08 | 51.84 | 10.08 |
| $\alpha$ | 412 | 424 | 65.79 | 2.63 | 61.84 | 2.63 |

The time-cluster with the peak ROC-AUC score for the α band in the young group showed a pattern in the left hemisphere with activity predicting incorrect morphosyntax in BA 22, BA 39, BA 40, and BA 42; BA 20, BA 22 and BA 40, and BA 37. In the right hemisphere, we found patterns predicting the incorrect morphosyntax in BA 22 and BA 7, and а pattern predicting correct morphosyntax in BA 22 and BA 39. For the high-ɣ, we found a smaller cluster in the left-hemispheric TPJ (BA 39) predicting correct morphosyntax, and some smaller left-hemispheric clusters in BA 1-2 and 4 predicting incorrect morphosyntax. In the right hemisphere, we found high-ɣ clusters in BA 7 predicting incorrect morphosyntax and BA 44 predicting correct morphosyntax.

For the older group, the peak ROC-AUC score showed a cluster in the left hemisphere predicting correct morphosyntax distributed across the BA 10, BA 39, BA 44, and BA 46. In the right hemisphere, there was a cluster predicting incorrect morphosyntax in the BA 39 and BA19.

## Discussion

We aimed at investigating the brain’s automatic language comprehension processes in healthy individuals of different ages at lexical, semantic, and morphosyntactic levels by scrutinising oscillatory brain activity in different frequency bands elicited by spoken words. By using beamformer-based source reconstruction and calculating ITPC for each point in source space for each word independently, we first combined the different words in into contrasts, one for each linguistic level (i.e., lexical, semantic, and morphosyntactic). We then proceeded to classify these contrasts in each of the five pre-defined frequency bands for both the young and the older group. We could, in line with previous research (M. Jensen et al., 2019), classify the different language processes, and found different time courses across the frequency bands. Crucially, the classification features differed not only between the bands and linguistic contrasts but also between the age groups.

### Lexical contrast

For the lexical condition, we found that the younger group had the fastest response differentiating meaningful vs. meaningless items already at ∼50 ms after the divergence point. This is line with the early lexical effects reported previously for unattended spoken words and pseudo-words (MacGregor et al., 2012; Shtyrov & Lenzen, 2017). Importantly, expanding on previous findings based on ERPs/ERFs, we found these lexicality-sensitive effects in oscillatory activity. Previously, the *γ* band activity has been linked to lexical processing (Towle et al., 2008) which aligns with our findings of significant clusters in the *γ* bands: for the young group in the medium-*γ* and high-*γ* band within the first 100 ms after the divergence point, and for the older group in the medium-*γ* band but only much later, at nearly 300 ms. At lower frequencies, the younger group had two clusters in the α band (at ∼230-260 ms), while the older group showed significant effects in the β band (at 266-280 ms). The latter finding fits with the suggestion that an increase in β band activity is related to cross-hemispheric activity (Schafer et al., 2014) that is involved in ageing-related compensation (Agarwal et al., 2016; Blasi et al., 2002; Manenti et al., 2013). That there was no high-*γ* activity for the older group may be due to a general gradual decrease in *γ* peak with age (Gaetz et al., 2012). The exact difference regarding the temporal dynamics of speech processing is hard to interpret from this data alone but a cautious proposition could be that more cross-hemispheric activity needs more time to transmit and synchronise information, which may contribute to the much later (by over 200 ms) onset of significant effects in the older group. Whether the lack of an early *γ* response in the older group is due to its actual absence or a larger temporal variance across participants with age is hard to tell from our data. Neuroanatomically, we found the lexical contrast to show bilateral patterns, involving the temporo-frontal language system as well as other, more distributed areas (note that, when interpreting these patterns, it is important to keep in mind that they demonstrate the most successful classification which does not necessarily correspond to the strongest activity in absolute terms).

### Semantic condition

In the semantic condition, we found patterns diverging in both time and frequency between the younger and older group, as in the lexical condition. The earliest successful decoding was for the young group in the β band at ∼142 ms, as opposed to 154 ms for the older group in the α band. The highest ROC-AUC score was for the young group and was also in higher frequency and earlier (∼150 ms; β) than in the older group (∼200 ms; α). Furthermore, the patterns from the classifiers also differed in other aspects. For the young group, most of the features in the β band predicted the action verb with activity in both hemispheres in both precentral motor (BAs 4 and 6) and inferior-frontal language (BA 44) areas. In contrast, the patterns of the older group were first predictive of the object noun in the α band, and only much later of the action verb in the medium-*γ* band. Furthermore, the activity used for the prediction of the action verb was mainly found in the right hemisphere for the older group, which implies a cross-hemispheric compensation involving cortical redistribution of semantic information processing.

While the young group’s decoding patterns were mostly expressed in the β band, the patterns for the older group were expressed in the α range instead. The finding of semantic activity in the β band for action verbs in the young group is in line with previously reported findings for word processing in general (Bastiaansen & Hagoort, 2006; M. Jensen et al., 2019). Furthermore, β band activity in the motor system has been linked to action control as well as action word processing in particular (O. Jensen et al., 2014; Vukovic & Shtyrov, 2014). Deterioration of the motor system with age is a well-established phenomenon (Seidler et al., 2010), which may explain the lack of similar β band activity in the older group. Instead, the classification was more successful for the visually-related object noun in the α band, which is known to be involved in visual information processing (e.g. O. Jensen & Mazaheri, 2010), and involved occipitotemporal areas (BA37) as well as classical language-processing areas in the temporal lobe, including the temporal pole which is often claimed to be the hub of lexical semantics (Lambon Ralph & Patterson, 2008). One might also speculate that the lower-frequency findings in the older group could be due to an age-related decrease in peak frequency, as α and β ranges are close to each other; however, the opposite (an increase in β with age) has also been reported previously (Gaetz et al., 2010; Ziegler et al., 2010), which makes the above neuroanatomically-based explanation more plausible. Finally, it could also represent an increased inter-regional amplitude correlation that was strongest in the α band range (Schafer et al., 2014). Further research is needed to address this observed shift in time and frequency of these semantic effects, as well as their neuroanatomical underpinnings.

In addition to these lower frequencies, both groups also exhibited medium-*γ* responses, which have been linked to semantic processing in previous studies (Bastiaansen & Hagoort, 2006; Lam et al., 2016; Levy et al., 2014). Interestingly, Wang et al. (2012) reported an increase in *γ* power tied to the predictability of a word rather than semantic integration; which may help explain the relatively modest *γ*-band effect here, given that we have a randomised stimulus presentation design with very low predictability of the stimuli.

### Morphosyntactic condition

For the morphosyntactic condition, we were able to successfully classify correct vs. incorrect morphosyntax for both groups. The findings in the two groups again diverged, although with a different pattern from the previous two contrasts, and thus in line with the general concept of syntax and lexical semantics being underpinned by distinct neurolinguistic systems (Friederici, 2002, 2012; Shtyrov, 2010). Firstly, both the young and the older groups showed activity in the high-*γ* band. Sentence-level syntactic binding has previously been reported to be linked β band synchrony (Wang, Zhu, et al., 2012) and low *γ* band activity (Weiss et al., 2005) with β and *γ* activity putatively related to working memory (Lundqvist et al., 2016, 2018). The high-*γ* band we found here, however, cannot be related to sentence-level binding and integration or working memory due to the morphosyntactic nature of the stimuli and unattended nature of the paradigm. Hence, the *γ* activity we found most likely was related to the processing of correct vs incorrect morphosyntax as such. Secondly, regarding the lower frequency bands, while the older group exhibited β band effects, the younger participants showed α activity. These findings of preserved *γ* activity across ages and β rather than α patterns in the older group are interesting as they suggest that the decrease in *γ* activity may not only be due to a decrease in physiological capacity but also (or even instead) reflect a change that is functionally specific to particular neurocognitive processes, which again supports the notion of distinct syntactic vs. lexical systems (Friederici & Weissenborn, 2007; Ullman, 2001). Regarding the timings of these effects, the young group showed them at ∼350 ms and ∼400 ms, i.e. exactly when syntactically related phenomena such as left-anterior negativity (LAN) are typically reported, including those reflecting syntactic agreement (De Vincenzi et al., 2003). Somewhat surprisingly, effects for the older group started earlier and extended for longer, with clusters ranging from 104 to 440 ms after the divergence point.

In the ERP/ERF literature, syntactic effects are typically reported in the left-lateralized core language areas, most commonly showing larger responses for grammatical anomalies in left inferior-frontal gyrus as well as in the left superior temporal areas (Hanna et al., 2014; Herrmann et al., 2011; Pulvermüller et al., 2005). In our study, the ITPC classification patterns were in line with those previous results, though somewhat more complex: left temporal and temporo-parietal areas predicted incorrect morphosyntax in the younger subjects alongside right-hemispheric activity (including right IFG for correct morphosyntax). For the older group, we found a very different pattern: temporo-parietal and frontal activity in the left hemisphere predicted correct morphosyntax, whereas the right-hemispheric activity indicated incorrect morphosyntax. A cautious interpretation could be that the highly automatised syntactic processing (Hahne & Friederici, 1999; Pulvermüller et al., 2008) involves rather limited resources under normal conditions with an additional activity/effort required to handle syntactic anomalies in the left temporo-frontal systems, which is what we observe in the younger subjects. With ageing, more resources are required, and, as compensation processes kick in, right-hemispheric activation becomes necessary for processing syntactic anomalies, whereas even the well-formed items lead to extended activation in the left temporo-frontal systems.

## General discussion

Taking the three contrasts together, we found that the spatiotemporal dynamics diverged between the young and the older group. In accordance with previous studies (e.g. Bastiaansen & Hagoort, 2006; M. Jensen et al., 2019; Lam et al., 2016), we found a difference in the frequency bands for the different linguistic processes. Furthermore, there is little overlap between the frequency bands for the young and the older groups in each specific contrast. While this needs further investigation to be fully explained, some preliminary speculative interpretations can still be offered based on diverging functional properties of the different frequency bands.

Gamma activity has been linked to GABA-ergic systems in the brain (Buzsáki & Wang, 2012; O. Jensen et al., 2014). GABA levels are lower in elderly compared to young participants (Maes et al., 2018), so a change in activity in this band is to be expected. Furthermore, *γ* activity is related to local processing (Buzsáki & Schomburg, 2015; Buzsáki & Silva, 2012) so the changes in the *γ* bands can be a way to investigate the compensation that happens with age. In that regard, we found a difference between the young and the elderly group especially for the lexical condition, with the young group showing multiple clusters in the high-*γ* and medium-*γ* bands not present in the elderly group. Another reason why we do not see early *γ* responses in the elderly group could be that the decrease in *γ*-power (Gaetz et al., 2012) and lower GABA levels result in more variable and inconsistent high-*γ* and medium-*γ* responses, which in turn lead to statistically less robust responses in the elderly group.

In contrast to *γ*, we found activity in the β band for the older group in the morphosyntactic condition, not expressed for the young group. This activity appeared at the latencies where we found no medium-*γ* activity in the young group, so it is unlikely to represent just a linear drop in peak frequency with age. The increase in β-band contribution with age could be related to cross-hemispheric communication, which becomes more prominent with age in order to engage right-hemispheric compensation processes, although further studies are needed to confirm this. In turn, oscillatory activity in the α band has been connected to both inhibition (O. Jensen & Mazaheri, 2010) and inter-regional communication (Bonnefond et al., 2017) and, as we mentioned above, both of those mechanisms may be at play in the semantic condition, which also requires further investigation.

In sum, using source-space MVPA analysis of ITPC MEG data, we have shown a detailed and complex picture of the alpha, beta, and gamma activity involved in different neural processes taking place during spoken language perception. Without any a priori selected times and/or areas of interest, this approach allowed for a controlled exploratory whole brain analysis that could reveal the spatiotemporal dynamics of the diverse cognitive processes that underpin automatic speech comprehension. Furthermore, we used a passive paradigm where the participants did not need to respond or actively react to the stimuli. This methodology (i.e., both the paradigm and the analysis techniques) can thus be applied not only to healthy younger and older participants, but also to patient groups, whose conditions prevent them from full cooperation with more active tasks, in order to further our understanding of the neural dynamics underpinning speech comprehension in both health and disease.

## Acknowledgements

MJ was supported by a grant from the Carlsberg Foundation, grant number CF16-000. BUW was supported by the European Union’s Horizon 2020 Marie Sklodowska Curie Individual Fellowship (grant agreement 893912). YS was supported by Danish Council for Independent Research (DFF 6110-00486, project 23776), Basic Research Program of the NRU Higher School of Economics (HSE University) and Lundbeck Foundation (grants R140-2013-12951, R164-2013-15801). Center of Functionally Integrative Neuroscience - CFIN/MIND*Lab* for funding Open Access. We declare no conflict of interest.

## Footnotes

1 The number of features was selected through a grid search between 512, 1024, and 2000 features at a single time point

## References

1. Abrams, L., & Farrell, M. T. (2011). Language processing in normal aging. The Handbook of Psycholinguistic and Cognitive Processes: Perspectives in Communication Disorders, 49–73.

2. Agarwal, S., Stamatakis, E. A., Geva, S., & Warburton, E. A. (2016). Dominant hemisphere functional networks compensate for structural connectivity loss to preserve phonological retrieval with aging. Brain and Behavior, 6(9), e00495. https://doi.org/10.1002/brb3.495

3. Bastiaansen, M., & Hagoort, P. (2006). Oscillatory neuronal dynamics during language comprehension. In C. Neuper & W. Klimesch (Eds.), Event-Related Dynamics of Brain Oscillations (Vol. 159, pp. 179–196). Elsevier. http://www.sciencedirect.com/science/article/pii/S0079612306590120

4. Blasi, V., Young, A. C., Tansy, A. P., Petersen, S. E., Snyder, A. Z., & Corbetta, M. (2002). Word Retrieval Learning Modulates Right Frontal Cortex in Patients with Left Frontal Damage. Neuron, 36(1), 159–170. https://doi.org/10.1016/S0896-6273(02)00936-4

5. Bonnefond, M., Kastner, S., & Jensen, O. (2017). Communication between Brain Areas Based on Nested Oscillations. ENeuro, 4(2). https://doi.org/10.1523/ENEURO.0153-16.2017

6. Buzsáki, G., & Schomburg, E. W. (2015). What does gamma coherence tell us about inter-regional neural communication? Nature Neuroscience, 18, 484–489. https://doi.org/10.1038/nn.3952

7. Buzsáki, G., & Silva, F. L. (2012). High frequency oscillations in the intact brain. Prog Neurobiol, 98, 241–249. https://doi.org/10.1016/j.pneurobio.2012.02.004

8. Buzsáki, G., & Wang, X. J. (2012). Mechanisms of gamma oscillations. Annu Rev Neurosci, 35, 203–225. https://doi.org/10.1146/annurev-neuro-062111-150444

9. Cohen, M. X. (2014). Analyzing neural time series data: Theory and practice. MIT Press.

10. Dalal, S. S., Vidal, J. R., Hamamé, C. M., Ossandón, T., Bertrand, O., Lachaux, J.-P., & Jerbi, K. (2011). Spanning the rich spectrum of the human brain: Slow waves to gamma and beyond. Brain Structure and Function, 216(2), 77–84. https://doi.org/10.1007/s00429-011-0307-z

11. Dale, A. M., Fischl, B., & Sereno, M. I. (1999). Cortical surface-based analysis. I. Segmentation and surface reconstruction. NeuroImage, 9, 179–194. https://doi.org/10.1006/nimg.1998.0395

12. Davis, S. W., Dennis, N. A., Buchler, N. G., White, L. E., Madden, D. J., & Cabeza, R. (2009). Assessing the effects of age on long white matter tracts using diffusion tensor tractography. NeuroImage, 46(2), 530–541. https://doi.org/10.1016/j.neuroimage.2009.01.068

13. De Vincenzi, M., Job, R., Di Matteo, R., Angrilli, A., Penolazzi, B., Ciccarelli, L., & Vespignani, F. (2003). Differences in the perception and time course of syntactic and semantic violations. Brain and Language, 85(2), 280–296.

14. Edmonds, B. A., & Krumbholz, K. (2014). Are interaural time and level differences represented by independent or integrated codes in the human auditory cortex? J Assoc Res Otolaryngol, 15(1), 103–114. https://doi.org/10.1007/s10162-013-0421-0

15. Friederici, A. D. (2002). Towards a neural basis of auditory sentence processing. Trends in Cognitive Sciences, 6(2), 78–84.

16. Friederici, A. D. (2012). The cortical language circuit: From auditory perception to sentence comprehension. Trends in Cognitive Sciences, 16(5), 262–268. https://doi.org/10.1016/j.tics.2012.04.001

17. Friederici, A. D., & Weissenborn, J. (2007). Mapping sentence form onto meaning: The syntax–semantic interface. Brain Research, 1146, 50–58.

18. Gaetz, W., Macdonald, M., Cheyne, D., & Snead, O. C. (2010). Neuromagnetic imaging of movement-related cortical oscillations in children and adults: Age predicts post-movement beta rebound. Neuroimage, 51(2), 792–807. https://doi.org/10.1016/j.neuroimage.2010.01.077

19. Gaetz, W., Roberts, T. P., Singh, K. D., & Muthukumaraswamy, S. D. (2012). Functional and structural correlates of the aging brain: Relating visual cortex (V1) gamma band responses to age-related structural change. Hum Brain Mapp, 33(9), 2035–2046. https://doi.org/10.1002/hbm.21339

20. Gansonre, C., Højlund, A., Leminen, A., Bailey, C., & Shtyrov, Y. (2018). Task-free auditory EEG paradigm for probing multiple levels of speech processing in the brain. Psychophysiology, 55(11), e13216. https://doi.org/10.1111/psyp.13216

21. Gertel, V. H., Karimi, H., Dennis, N. A., Neely, K. A., & Diaz, M. T. (2020). Lexical frequency affects functional activation and accuracy in picture naming among older and younger adults. Psychology and Aging. https://doi.org/10.1037/pag0000454

22. Gramfort, A., Luessi, M., Larson, E., Engemann, D. A., Strohmeier, D., Brodbeck, C., Goj, R., Jas, M., Brooks, T., Parkkonen, L., & Hämäläinen, M. (2013). MEG and EEG data analysis with MNE-Python. Frontiers in Neuroscience, 7, 267. https://doi.org/10.3389/fnins.2013.00267

23. Hagoort, P., & Brown, C. M. (2000). ERP effects of listening to speech: Semantic ERP effects. Neuropsychologia, 38(11), 1518–1530. https://doi.org/10.1016/S0028-3932(00)00052-X

24. Hahne, A., & Friederici, A. D. (1999). Electrophysiological evidence for two steps in syntactic analysis: Early automatic and late controlled processes. Journal of Cognitive Neuroscience, 11(2), 194–205.

25. Hanna, J., Mejias, S., Schelstraete, M.-A., Pulvermüller, F., Shtyrov, Y., & van der Lely, H. K. J. (2014). Early activation of Broca’s area in grammar processing as revealed by the syntactic mismatch negativity and distributed source analysis. Cognitive Neuroscience, 5(2), 66–76. https://doi.org/10.1080/17588928.2013.860087

26. Hari, R., Joutsiniemi, S. L., & Sarvas, J. (1988). Spatial resolution of neuromagnetic records: Theoretical calculations in a spherical model. Electroencephalogr Clin Neurophysiol, 71(1), 64–72.

27. Herrmann, B., Maess, B., & Friederici, A. D. (2011). Violation of syntax and prosody— Disentangling their contributions to the early left anterior negativity (ELAN). Neuroscience Letters, 490(2), 116–120. https://doi.org/10.1016/j.neulet.2010.12.039

28. Hyder, R., Højlund, A., Jensen, M., Johnsen, E. L., Østergaard, K., & Shtyrov, Y. (2021). STN-DBS affects language processing differentially in Parkinson’s disease: Multiple-case MEG study. Acta Neurologica Scandinavica, 144(2), 132–141. https://doi.org/10.1111/ane.13423

29. Hyder, R., Højlund, A., Jensen, M., Østergaard, K., & Shtyrov, Y. (2020). Objective assessment of automatic language comprehension mechanisms in the brain: Novel E/MEG paradigm. Psychophysiology, 57(5). https://doi.org/10.1111/psyp.13543

30. Jas, M., Engemann, D. A., Bekhti, Y., Raimondo, F., & Gramfort, A. (2017). Autoreject: Automated artifact rejection for MEG and EEG data. NeuroImage, 159, 417–429. https://doi.org/10.1016/j.neuroimage.2017.06.030

31. Jensen, M., Hyder, R., & Shtyrov, Y. (2019). MVPA analysis of intertrial phase coherence of neuromagnetic responses to words reliably classifies multiple levels of language processing in the brain. ENeuro. https://doi.org/10.1523/ENEURO.0444-18.2019

32. Jensen, O., & Mazaheri, A. (2010). Shaping Functional Architecture by Oscillatory Alpha Activity: Gating by Inhibition. Frontiers in Human Neuroscience, 4. https://doi.org/10.3389/fnhum.2010.00186

33. Jensen, O., Spaak, E., & Zumer, J. M. (2014). Human brain oscillations: From physiological mechanisms to analysis and cognition. In S. Supek & C. J. Aine (Eds.), Magnetoencephalography: From signals to dynamic cortical networks (pp. 359–403). Springer Berlin Heidelberg.

34. Kielar, A., Deschamps, T., Jokel, R., & Meltzer, J. A. (2016). Functional reorganization of language networks for semantics and syntax in chronic stroke: Evidence from MEG: Reorganization of Language Networks in Chronic Stroke. Human Brain Mapping, 37(8), 2869–2893. https://doi.org/10.1002/hbm.23212

35. Kösem, A., & van Wassenhove, V. (2017). Distinct contributions of low- and high-frequency neural oscillations to speech comprehension. Language, Cognition and Neuroscience, 32(5), 536–544. https://doi.org/10.1080/23273798.2016.1238495

36. Lam, N. H. L., Schoffelen, J.-M., Uddén, J., Hultén, A., & Hagoort, P. (2016). Neural activity during sentence processing as reflected in theta, alpha, beta, and gamma oscillations. NeuroImage, 142, 43–54. https://doi.org/10.1016/j.neuroimage.2016.03.007

37. Lambon Ralph, M. A., & Patterson, K. (2008). Generalization and Differentiation in Semantic Memory: Insights from Semantic Dementia. Annals of the New York Academy of Sciences, 1124(1), 61–76. https://doi.org/10.1196/annals.1440.006

38. Levy, J., Hagoort, P., & Démonet, J.-F. (2014). A neuronal gamma oscillatory signature during morphological unification in the left occipitotemporal junction. Human Brain Mapping, 35(12), 5847–5860. https://doi.org/10.1002/hbm.22589

39. Lundqvist, M., Herman, P., Warden, M. R., Brincat, S. L., & Miller, E. K. (2018). Gamma and beta bursts during working memory readout suggest roles in its volitional control. Nat Commun, 9(1), 394. https://doi.org/10.1038/s41467-017-02791-8

40. Lundqvist, M., Rose, J., Herman, P., Brincat, S. L., Buschman, T. J., & Miller, E. K. (2016). Gamma and Beta Bursts Underlie Working Memory. Neuron, 90(1), 152–164. https://doi.org/10.1016/j.neuron.2016.02.028

41. Luo, H., & Poeppel, D. (2007). Phase Patterns of Neuronal Responses Reliably Discriminate Speech in Human Auditory Cortex. Neuron, 54(6), 1001–1010. https://doi.org/10.1016/j.neuron.2007.06.004

42. MacGregor, L., Pulvermüller, F., van Casteren, M., & Shtyrov, Y. (2012). Ultra-rapid access to words in the brain. Nature Communications, 3(1). https://doi.org/10.1038/ncomms1715

43. Maes, C., Hermans, L., Pauwels, L., Chalavi, S., Leunissen, I., Levin, O., Cuypers, K., Peeters, R., Sunaert, S., Mantini, D., Puts, N. A. J., Edden, R. A. E., & Swinnen, S. P. (2018). Age-related differences in GABA levels are driven by bulk tissue changes. Human Brain Mapping, 39(9), 3652–3662. https://doi.org/10.1002/hbm.24201

44. Manenti, R., Brambilla, M., Petesi, M., Miniussi, C., & Cotelli, M. (2013). Compensatory networks to counteract the effects of ageing on language. Behavioural Brain Research, 249, 22–27. https://doi.org/10.1016/j.bbr.2013.04.011

45. Naatanen, R., Paavilainen, P., Rinne, T., & Alho, K. (2007). The mismatch negativity (MMN) in basic research of central auditory processing: A review. Clin Neurophysiol, 118(12), 2544–2590. https://doi.org/10.1016/j.clinph.2007.04.026

46. Oldfield, R. C. (1971). The assessment and analysis of handedness: The Edinburgh inventory. Neuropsychologia, 9(1), 97–113. https://doi.org/10.1016/0028-3932(71)90067-4

47. Pedregosa, F., Varoquaux, G., Gramfort, A., Michel, V., Thirion, B., Grisel, O., Blondel, M., Prettenhofer, P., Weiss, R., Dubourg, V., Vanderplas, J., Passos, A., Cournapeau, D., Brucher, M., Perrot, M., & Duchesnay, E. (2011). Scikit-learn: Machine Learning in Python. Journal of Machine Learning Research, 12, 2825–2830.

48. Pfefferbaum, A., Sullivan, E. V., Hedehus, M., Lim, K. O., Adalsteinsson, E., & Moseley, M. (2000). Age-related decline in brain white matter anisotropy measured with spatially corrected echo-planar diffusion tensor imaging. Magnetic Resonance in Medicine, 44(2), 259–268.

49. Pulvermüller, F. (2005). Brain mechanisms linking language and action. Nature Reviews Neuroscience, 6(7), 576–582. https://doi.org/10.1038/nrn1706

50. Pulvermüller, F., & Fadiga, L. (2010). Active perception: Sensorimotor circuits as a cortical basis for language. Nature Reviews Neuroscience, 11(5), 351–360. https://doi.org/10.1038/nrn2811

51. Pulvermüller, F., Huss, M., Kherif, F., Moscoso del Prado Martin, F., Hauk, O., & Shtyrov, Y. (2006). Motor cortex maps articulatory features of speech sounds. Proceedings of the National Academy of Sciences, 103(20), 7865–7870. https://doi.org/10.1073/pnas.0509989103

52. Pulvermuller, F., & Shtyrov, Y. (2006). Language outside the focus of attention: The mismatch negativity as a tool for studying higher cognitive processes. Prog Neurobiol, 79(1), 49–71. https://doi.org/10.1016/j.pneurobio.2006.04.004

53. Pulvermüller, F., Shtyrov, Y., Hasting, A. S., & Carlyon, R. P. (2008). Syntax as a reflex: Neurophysiological evidence for early automaticity of grammatical processing. Brain and Language, 104(3), 244–253.

54. Pulvermüller, F., Shtyrov, Y., & Ilmoniemi, R. (2005). Brain Signatures of Meaning Access in Action Word Recognition. Journal of Cognitive Neuroscience, 17(6), 884–892. https://doi.org/10.1162/0898929054021111

55. Raz, N., Lindenberger, U., Rodrigue, K. M., Kennedy, K. M., Head, D., Williamson, A., Dahle, C., Gerstorf, D., & Acker, J. D. (2005). Regional Brain Changes in Aging Healthy Adults: General Trends, Individual Differences and Modifiers. Cerebral Cortex, 15(11), 1676–1689. https://doi.org/10.1093/cercor/bhi044

56. Schafer, C. B., Morgan, B. R., Ye, A. X., Taylor, M. J., & Doesburg, S. M. (2014). Oscillations, networks, and their development: MEG connectivity changes with age. Hum Brain Mapp, 35(10), 5249–5261. https://doi.org/10.1002/hbm.22547

57. Seidler, R. D., Bernard, J. A., Burutolu, T. B., Fling, B. W., Gordon, M. T., Gwin, J. T., Kwak, Y., & Lipps, D. B. (2010). Motor control and aging: Links to age-related brain structural, functional, and biochemical effects. Neuroscience & Biobehavioral Reviews, 34(5), 721–733. https://doi.org/10.1016/j.neubiorev.2009.10.005

58. Sekihara, K., & Nagarajan, S. S. (2008). Adaptive Spatial Filters for Electromagnetic Brain Imaging. Springer Berlin Heidelberg. http://link.springer.com/10.1007/978-3-540-79370-0

59. Shtyrov, Y. (2010). Automaticity and attentional control in spoken language processing: Neurophysiological evidence. The Mental Lexicon, 5(2), 255–276. https://doi.org/10.1075/ml.5.2.06sht

60. Shtyrov, Y., & Lenzen, M. (2017). First-pass neocortical processing of spoken language takes only 30 msec: Electrophysiological evidence. Cognitive Neuroscience, 8(1), 24– 38. https://doi.org/10.1080/17588928.2016.1156663

61. Teng, X., Tian, X., Rowland, J., & Poeppel, D. (2017). Concurrent temporal channels for auditory processing: Oscillatory neural entrainment reveals segregation of function at different scales. PLOS Biology, 15(11), e2000812. https://doi.org/10.1371/journal.pbio.2000812

62. Thiel, A., Habedank, B., Herholz, K., Kessler, J., Winhuisen, L., Haupt, W. F., & Heiss, W.-D. (2006). From the left to the right: How the brain compensates progressive loss of language function. Brain and Language, 98(1), 57–65. https://doi.org/10.1016/j.bandl.2006.01.007

63. Thielscher, A., Antunes, A., & Saturnino, G. B. (2015). Field modeling for transcranial magnetic stimulation: A useful tool to understand the physiological effects of TMS? Engineering in Medicine and Biology Society (EMBC), 2015 37th Annual International Conference of the IEEE, 222–225. http://ieeexplore.ieee.org/xpls/abs_all.jsp?arnumber=7318340

64. Towle, V. L., Yoon, H. A., Castelle, M., Edgar, J. C., Biassou, N. M., Frim, D. M., Spire, J. P., & Kohrman, M. H. (2008). ECoG gamma activity during a language task: Differentiating expressive and receptive speech areas. Brain, 131(8), 2013–2027. https://doi.org/10.1093/brain/awn147

65. Tyler, L. K., & Marslen-Wilson, W. (2008). Fronto-temporal brain systems supporting spoken language comprehension. Philosophical Transactions of the Royal Society B: Biological Sciences, 363(1493), 1037–1054. https://doi.org/10.1098/rstb.2007.2158

66. Tyler, L. K., Shafto, M. A., Randall, B., Wright, P., Marslen-Wilson, W. D., & Stamatakis, E. A. (2010). Preserving Syntactic Processing across the Adult Life Span: The Modulation of the Frontotemporal Language System in the Context of Age-Related Atrophy. Cerebral Cortex, 20(2), 352–364. https://doi.org/10.1093/cercor/bhp105

67. Tyler, L. K., Wright, P., Randall, B., Marslen-Wilson, W. D., & Stamatakis, E. A. (2010). Reorganization of syntactic processing following left-hemisphere brain damage: Does right-hemisphere activity preserve function? Brain, 133(11), 3396–3408. https://doi.org/10.1093/brain/awq262

68. Ullman, M. T. (2001). The declarative/procedural model of lexicon and grammar. Journal of Psycholinguistic Research, 30(1), 37–69.

69. Van Veen, B. D., Van Drongelen, W., Yuchtman, M., & Suzuki, A. (1997). Localization of brain electrical activity via linearly constrained minimum variance spatial filtering. IEEE Transactions on Biomedical Engineering, 44(9), 867–880. https://doi.org/10.1109/10.623056

70. Varoquaux, G., Raamana, P. R., Engemann, D. A., Hoyos-Idrobo, A., Schwartz, Y., & Thirion, B. (2017). Assessing and tuning brain decoders: Cross-validation, caveats, and guidelines. NeuroImage, 145, 166–179. https://doi.org/10.1016/j.neuroimage.2016.10.038

71. Vukovic, N., & Shtyrov, Y. (2014). Cortical motor systems are involved in second-language comprehension: Evidence from rapid mu-rhythm desynchronisation. Neuroimage, 102 *Pt* *2*, 695–703. https://doi.org/10.1016/j.neuroimage.2014.08.039

72. Wang, L., Jensen, O., van den Brink, D., Weder, N., Schoffelen, J. M., Magyari, L., Hagoort, P., & Bastiaansen, M. (2012). Beta oscillations relate to the N400m during language comprehension. Hum Brain Mapp, 33(12), 2898–2912. https://doi.org/10.1002/hbm.21410

73. Wang, L., Zhu, Z., & Bastiaansen, M. (2012). Integration or Predictability? A Further Specification of the Functional Role of Gamma Oscillations in Language Comprehension. Frontiers in Psychology, 3. https://doi.org/10.3389/fpsyg.2012.00187

74. Weiss, S., Mueller, H., Schack, B., King, J., Kutas, M., & Rappelsberger, P. (2005). Increased neuronal communication accompanying sentence comprehension. International Journal of Psychophysiology, 57(2), 129–141. https://doi.org/10.1016/j.ijpsycho.2005.03.013

75. Westner, B. (2017). The Hilbert beamformer pipeline. https://brittas-summerofcode.blogspot.com/2017/08/the-hilbert-beamformer-pipeline_29.html

76. Westner, B., & Dalal, S. S. (2017). Faster than the brain’s speed of light: Retinocortical interactions differ in high frequency activity when processing darks and lights. BioRxiv. https://doi.org/10.1101/153551

77. Ziegler, D. A., Pritchett, D. L., Hosseini-Varnamkhasti, P., Corkin, S., Hamalainen, M., Moore, C. I., & Jones, S. R. (2010). Transformations in oscillatory activity and evoked responses in primary somatosensory cortex in middle age: A combined computational neural modeling and MEG study. Neuroimage, 52(3), 897–912. https://doi.org/10.1016/j.neuroimage.2010.02.004

